# How random is the review outcome? A systematic study of the impact of external factors on *eLife* peer review

**DOI:** 10.1101/2023.01.04.522708

**Authors:** Weixin Liang, Kyle Mahowald, Jennifer Raymond, Vamshi Krishna, Daniel Smith, Daniel Jurafsky, Daniel McFarland, James Zou

## Abstract

The advance of science rests on a robust peer review process. However whether or not a paper is accepted can depend on random external factors--e.g. the timing of the submission, the matching of editors and reviewers--that are beyond the quality of the work. This article systematically investigates the impact of these random factors independent of the paper’s quality on peer review outcomes in a major biomedical journal, *eLife*. We analyzed all of the submissions to *eLife* between 2016 to 2018, with 23,190 total submissions. We examined the effects of random factors at each decision point in the review process, from the gatekeeping senior editors who may desk-reject papers to review editors and reviewers who recommend the final outcome. Our results suggest that the peer-review process in *eLife* is robust overall and that random external factors have relatively little quantifiable bias.

## Introduction

The establishment of scientific facts rests on a robust review process. Peer review is expected to follow the organizing ethos of being impartial, systematic, and universalistic (Merton, 1973), where each submission is judged solely on its intellectual merit. However, the scientific peer review process is often seen as opaque and inconsistent to authors (Rennie, 2016). For example, a major computer science conference (NeurIPS) performed an experiment where 10% of the submissions were independently reviewed by two sets of reviewers, and found that about 57% of the papers accepted by one committee were rejected by the other (Price, 2014). This suggests that external factors that are independent of the intrinsic quality of the work such as which editor/reviewers happen to be available, etc.--might substantially impact the review outcome in certain disciplines. Such external factors introduce noise into the peer review process which can frustrate researchers and slow the progress of science.

In this paper, we systematically quantify the extent to which factors external to a paper’s quality affect how likely the paper is accepted in a major biomedical journal, *eLife*. We focus on factors that have a large element of randomness, such as when the paper is submitted and the matching of editors and reviewers, which can depend on individual availabilities. We analyze each decision step of the review process from the gate-keeping senior editors who may desk-reject papers to review editors and reviewers who recommend the final outcome. To do so, we leverage the comprehensive data on the full review process of 34,161 *eLife* submissions. We find that the peer-review process in *eLife* is relatively robust and that the external factors examined here have little quantifiable effects on the paper outcomes.

Much of the previous research on peer review investigates the extent to which reviews are unbiased, efficient, and predictive (Bornmann, 2011; Marsh et al., 2008; Siler et al., 2015). Bias enters the peer review when extra-scientific factors — social characteristics and phenomena external to the science reported — play a role in explaining the desk decisions, reviewer recommendations, and the final outcome to publish (Zuckerman & Merton, 1971). Several previous works analyzed systematic bias toward underrepresented demographic groups, such as women and non-white, non-Anglo-American scientists (Bedi et al., 2012; Fox et al., 2016; Hopkins et al., 2013; Primack et al., 2009). While findings are mixed, a core insight from this research is that the alignment of statuses (i.e. homophily) and closer professional ties (i.e. nepotism) among the authors and reviewing scientists could result in unfair advantage (Demarest et al., 2014; Sandström & Hällsten, 2008; Teplitskiy et al., 2018; Wennerås & Wold, 1997). For example, a previous analysis of a smaller set of *eLife* submissions identified homophily between gatekeepers of the journal and authors (Murray et al., 2019). Yet it is often challenging to fully disentangle demographic factors from the type and quality of the research, especially when competent reviewers work adjacent to submitting authors within narrow research specializations (Li, 2017; Li & Agha, 2015; Mallard et al., 2009).

Still more, we know less about the effect of seemingly arbitrary factors that might play a role in the review process. For example, the assignment of reviewers often depends on which reviewers happened to be available, which can have some randomness to it. However, we don’t know how much this randomness systematically impacts the review outcomes in the life sciences. Some works suggest that increased demand to publish translates to heightened burdens on reviewers, resulting in more frequent assignments to less competent and more overworked reviewers (Arns, 2014; Kovanis et al., 2016). Other work shows a negative association between the number of submissions reviewed and harsher (lower) ratings (Jayasinghe et al., 2003; Marsh et al., 2008). While some studies focus on the timing of submissions with respect to important, policy-relevant dates in the organizational history of the respective granting and publishing institutions, we know of no studies that investigate the relationship of the quotidian timing submissions and their initial and final outcomes (Gallo et al., 2016). These contextual factors, while not likely driven by bias from social and intellectual prejudices, can nonetheless systematically introduce noise into the review process resulting in idiosyncratic, seemingly arbitrary outcomes.

This motivates us to focus on those features of the review process, which anecdotally can affect paper’s acceptance, but have been underexplored in the literature. In particular, we focus on studying two possible sources of arbitrariness: the timing of the submission and the assignment of editors and reviewers. Our study opens up a new perspective for analyzing the robustness of peer review by bringing into view two important contextual features of the process itself.

## Results

### Overview of the *eLife* submissions data

*eLife* is a major biomedical journal that follows a three-stage peer review process. As shown in Figure 1a, initial submissions are first assessed by a senior editor who may either desk-reject or encourage the authors to provide a full submission. Between 2012 and 2018, the majority, 72%, of initial submissions were desk-rejected at this stage. The remaining submissions underwent review by several reviewers, who provided independent evaluations, followed by a decision-making process by the reviewing editors. Although revisions may be requested at this stage, our study focuses on the final decision of the reviewing editors. We were granted comprehensive access to all aspects of the review process for each submission through an agreement with *eLife*, including original submissions and reviewer text. Further details on the dataset can be found in the Materials and Methods section.

**Figure 1:**
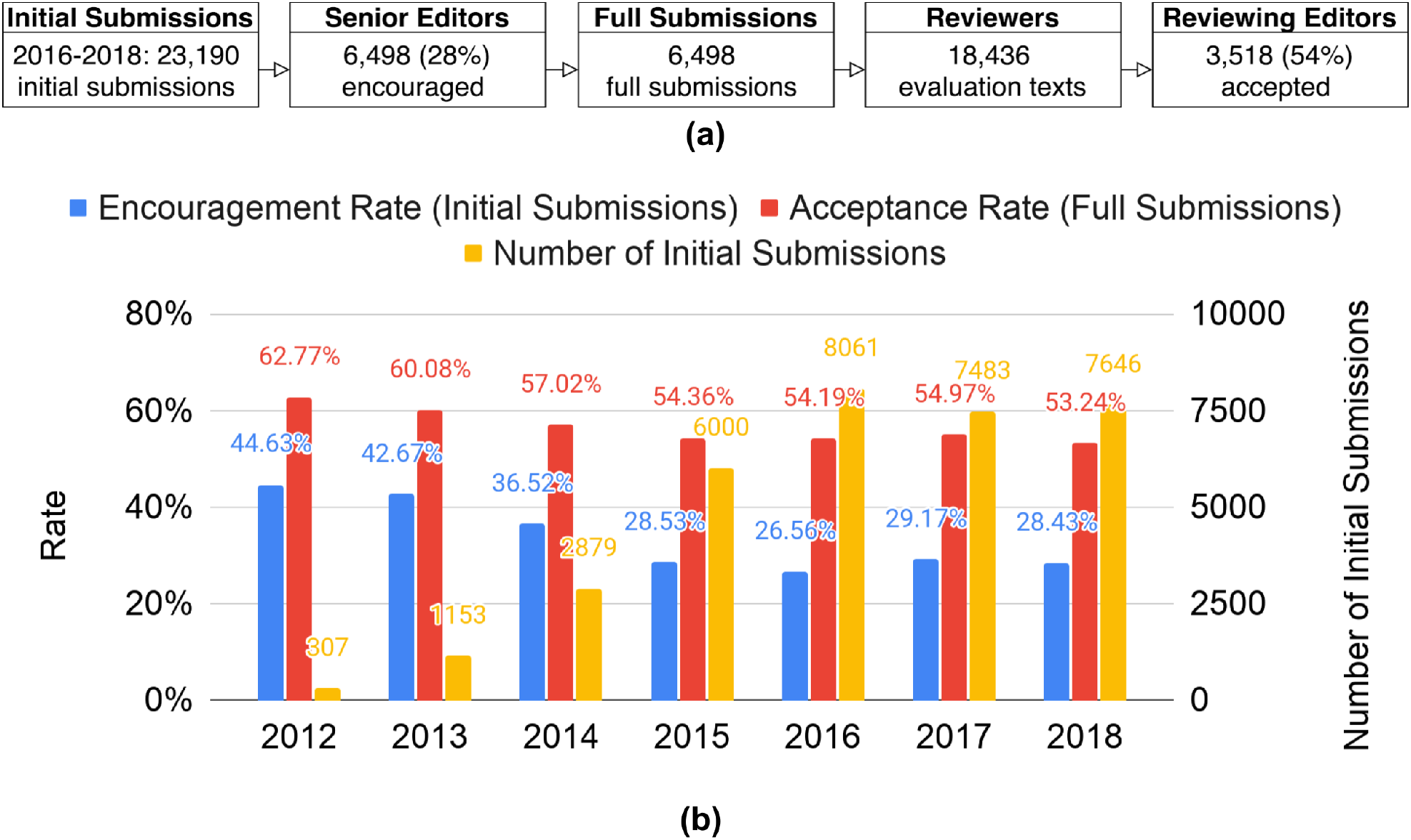
Overview of *eLife* review process and dataset. (a) Flow of submissions in the *eLife* review process. Initial submissions are assessed by senior editors, who may reject them outright. Full submissions are reviewed by reviewers and then recommended for acceptance or rejection by review editors. (b) Submissions and selectivity of *eLife* over the years. The x-axis indicates the year of submission. The orange bars represent the annual number of initial submissions, the blue bars represent the encouragement rate of initial submissions, and the red bar represents the acceptance rate of full submissions. The submission statistics stabilized in 2016-2018, which is why we focus on these years in our analysis to ensure the robustness of our findings.

Figure 1b presents the submissions and selectivity of *eLife* over the years. Our dataset includes the peer review outcomes of *eLife* from its founding in 2012 to 2018, encompassing a total of 34,161 initial submissions and 10,227 full submissions. To ensure the robustness of our findings, we focus specifically on the period between 2016 and 2018, during which the submission and selectivity statistics were relatively stable. This results in a study of 23,190 initial submissions and 6,498 full submissions to *eLife* submitted between 2016 and 2018.

In the subsequent sections, we present our analysis in a manner that reflects the three-stage review process of *eLife*. We begin by examining the impact of external factors on the decisions of senior editors, followed by an analysis of the influence of such factors on reviewers and reviewing editors. For each stage, we focus on two types of external factors that may vary: the timing of the submission and the assignment of editors and reviewers.

### Analysis of senior editor decisions

To begin, we examined whether the likelihood of a senior editor encouraging a submission is impacted by the number of submissions the editor is handling at a given time. Figure 2a shows that, despite fluctuations in the number of submissions throughout the year with peaks at the beginning of the year and in the summer, the encouragement rate remains constant. This suggests that the senior editors’ decisions are relatively unaffected by the external factor of the number of submissions, given that the quality of submissions is assumed to be consistent throughout the year.

**Figure 2:**
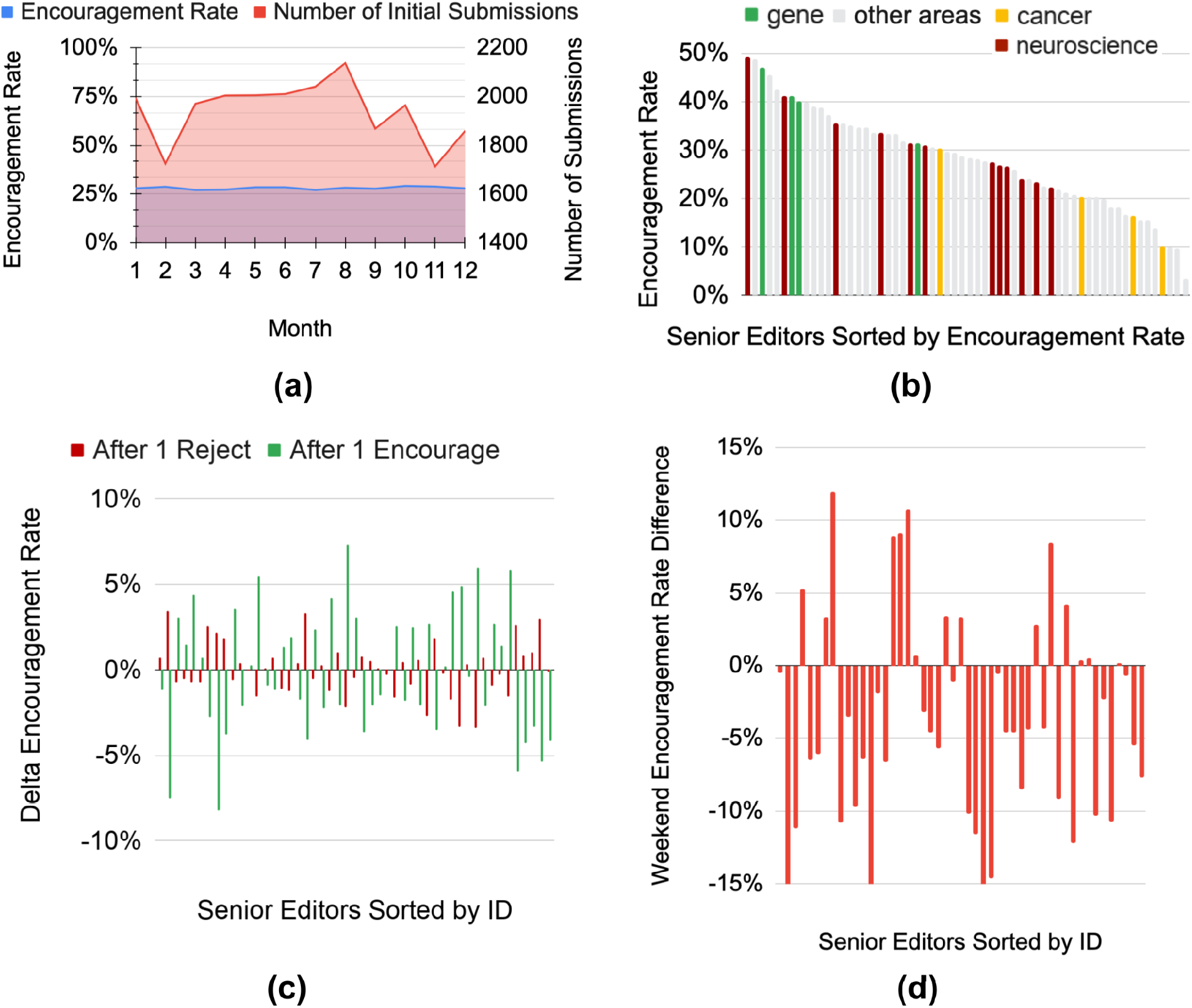
Analysis of senior editor decisions. (a) No busyness effect: The x-axis indicates the submission month, with the red curve representing the number of submissions and the blue curve representing the encouragement rate for that month. Although the number of submissions varies throughout the year, with peaks at the beginning of the calendar year and in the summer, the encouragement rate remains constant. (b) Heterogeneous encouragement rates: Each bar represents the encouragement rate of a senior editor. Red bars indicate senior editors in neuroscience, green bars indicate senior editors in gene expression, yellow bars indicate senior editors in cancer, and gray bars indicate senior editors in other disciplines. Senior editors are sorted by their encouragement rates for better visualization. The encouragement rates range from 9.9% to 49.7%. While different disciplines have different overall encouragement rates, there is also heterogeneity within the same discipline. (c) No anchoring effect: Each green bar, red bar pair represents a senior editor. The value of each green bar indicates the senior editor’s encouragement rate when the previous decision was an encouragement, while the value of each red bar indicates the senior editor’s encouragement rate when the previous decision was a rejection. There is no consistent trend, indicating that senior editors are robust to the anchoring effect. (d) Individual analysis of the weekend effect: Each red bar represents a senior editor. The value of each bar indicates the difference between the senior editor’s encouragement rate on weekends and on weekdays. Most of the bars are negative, indicating that most senior editors have a lower encouragement rate on weekends.

We next explored the potential impact of the arbitrary ordering of submissions on a paper’s outcome. Specifically, we investigated whether the last decision made by a senior editor (i.e. to encourage or desk-reject) influences the likelihood of the editor encouraging the subsequent submission. According to psychological studies (Tversky and Kahneman, 1974), the anchoring effect can lead to human judgments and evaluations being influenced by initial information - in this case, the previous paper - even when it is irrelevant. Figure 2c shows the conditional encouragement rates of each senior editor based on a previous rejection or a previous encouragement. There is no apparent trend (p=0.732), indicating that senior editors’ initial decisions to encourage or desk-reject are not influenced by the anchoring effect.

Surprisingly, the average encouragement rate for editorial decisions made on weekends (0.238 ± 0.013) was significantly lower than for decisions made on weekdays (0.290 ± 0.006) (χ^2^(df=1, n=23,190)=47.979, *p*=4.3075e-12), and this effect held true for the majority of senior editors (Figure 2d, Figure S2).

There are several possible explanations for this difference in encouragement rates on weekends. One possibility is that senior editors may delay reviewing lower-quality manuscripts or delivering negative news until the weekend. However, rejection tends to occur faster for senior editors compared to encouragement (6.09 days on average for rejection compared to 7.34 days for encouragement, as shown in Figure S3), which does not support this explanation. It is also possible that the mental states of senior editors who must work on weekends may differ due to their busy schedules. Further experiments are necessary to fully understand the cause of this weekend effect.

After examining the impact of submission timing, we next investigate the extent to which the assignment of submissions to senior editors could affect their outcomes. Figure 2b reveals that different senior editors had significantly different encouragement rates, ranging from 9.9% to 49.7%, with a 502% relative difference between the extremes. Encouragement rates also varied by research discipline, and even within the same discipline, there was heterogeneity. For example, among the 11 senior editors in neuroscience, encouragement rates ranged from 23% to 40%. Cancer research generally had lower encouragement rates than neuroscience, ranging from 10% to 30% (yellow bars in Figure 2b), a threefold difference. Some of the observed heterogeneity in acceptance rates may be due to differences in the selectivity of different senior editors. There may also be variations in the quality of submissions to *eLife* across different research disciplines. For instance, certain disciplines may have a greater number of other prestigious journals relative to *eLife*, which could lead to a lower quality of submissions to *eLife* and consequently a lower encouragement rate.

To gain further insight into the heterogeneity of senior editors, we analyzed their initial decisions to encourage or reject against the final acceptance rate of the encouraged papers, which serves as a proxy for paper quality. The results showed no correlation between each senior editor’s encouragement rate and the final acceptance rate (determined by the reviewing editors’ decisions) of the submissions they encouraged (Figure S1, r=0.015, p=0.903). One potential explanation for this lack of correlation is that senior editors are highly calibrated with one another. If the submissions that senior editors encourage after desk rejection have similar quality, this should generally lead to similar acceptance rates among the reviewing editors.

### Analysis of reviewer decisions

Following the senior editor’s decision to encourage a full submission, the next step in the manuscript evaluation process is review by several reviewers. As eLife reviews do not include an explicit review rating score, we performed sentiment analysis on each review text to quantify its positivity. The sentiment scores were generated using a deep neural language model that was calibrated to expert evaluations (see Methods). Since there are typically multiple reviews for each submission, we controlled for submission quality by examining whether some reviewers consistently have higher or lower review ratings compared to other reviewers of the same submissions.

To better understand the level of bias present in the decisions made by reviewers, we compared the actual distribution of average normalized review ranks for each reviewer with three simulated distributions representing different levels of bias through a simulation study. To ensure statistical power, we focused on reviewers with at least 10 reviews. The real distribution was based on our sentiment analysis of the review texts, and represented the actual ratings given by reviewers to submissions. The simulated distribution with 0% bias represented a scenario of fair peer review, in which all reviewers had an equal likelihood of giving a high or low review score. This was achieved by drawing random ratings from the standard Gaussian distribution N(0, 1), which was independent of the actual ratings given by reviewers. The simulated distribution with 5% bias represented systematically biased ratings, with 2.5% of reviewers consistently giving higher ratings and another 2.5% consistently giving lower ratings. These biased ratings were drawn from the Gaussian distributions N(1,1) and N(−1,1) respectively. As shown in Figure 3a, the yellow curve (simulation with 5% bias) closely matches the blue curve (real distribution). This suggests that the level of systematic bias in the peer review process is relatively low.

**Figure 3:**
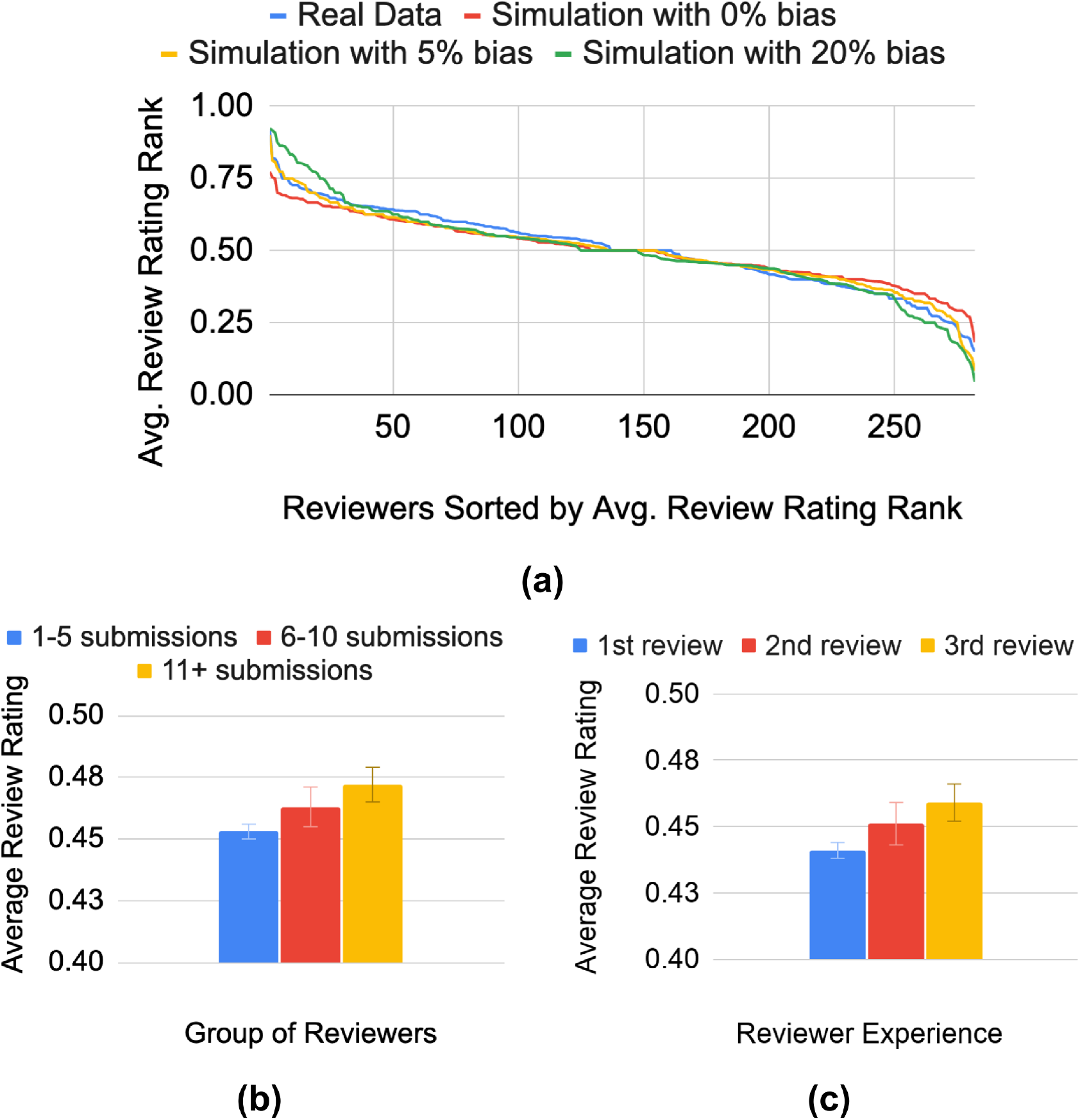
Analysis of reviewer decisions. (a) Heterogeneity of reviewers: The y-axis represents the average review rating rank, with a rank of 1.0 indicating that the reviewer consistently gives the most positive review compared to other reviewers for the same submission. The blue curve shows the actual distribution of average review rating ranks among reviewers. The red, yellow, and green curves represent simulations with 0%, 5%, and 20% bias, respectively. The yellow curve (simulation with 5% bias) aligns with the blue curve (real data). (b) New reviewer effect: The y-axis represents the average review rating. The blue bar shows the average review rating for reviewers who have completed 1-5 *eLife* reviews, the red bar shows the average review rating for reviewers who have completed 6-10 *eLife* reviews, and the green bar shows the average review rating for reviewers who have completed 11 or more *eLife* reviews. More experienced reviewers tend to give more positive reviews. (c) For reviewers who have completed at least three *eLife* reviews, the blue bar shows the average review rating for their first *eLife* review, the red bar shows the average review rating for their second *eLife* review, and the orange bar shows the average review rating for their third *eLife* review. As reviewers gain more experience with *eLife*, they give more positive reviews on average.

In addition, we included a simulation with 20% bias as a power analysis to demonstrate the sensitivity of our method. If 20% of reviewers were systematically biased as in our model, we would have detected a notable difference in the curves. The fact that we did not observe such a difference further supports the conclusion that the level of systematic bias is relatively low. It is worth noting that there could be other bias models to explore in future work. For example, biased reviewers could be two standard deviations off (i.e., N(2,1), N(−2,1)) rather than one standard deviation (i.e., N(1,1), N(−1,1)). Overall, our simulation study provides valuable insights into the level of bias present in the peer review process. While we have found that the level of bias is relatively low, it is important to continue monitoring and addressing any potential biases in order to ensure the fairness and integrity of our review process.

Furthermore, our analysis did not reveal any significant differences in the average review ratings based on the gender or nationality of the reviewers. When comparing U.S. reviewers (0.457 ± 0.004) to non-U.S. reviewers (0.456 ± 0.003), we found that there was no significant difference, as indicated by a χ2 test with df=1, n=18,434, and p=0.904. Similarly, when comparing male reviewers (0.459 ± 0.003) to female reviewers (0.452 ± 0.006), we found no significant difference, with a χ2 test yielding df=1, n=18,434, and p=0.478. These results suggest that reviewer gender and nationality do not significantly impact review ratings in the peer review process.

### New reviewer effect

In addition to studying experienced reviewers, we also examined the ratings of less experienced reviewers. Our findings showed that reviewers with less experience with eLife gave lower review ratings on average. As illustrated in Figure 3b, the average review rating for reviewers who had reviewed 1-5 eLife submissions was 0.453 ± 0.003, while the average rating for reviewers who had reviewed 5-10 submissions was 0.463 ± 0.008. For reviewers who had reviewed 10 or more submissions, the average rating was 0.472 ± 0.007. This demonstrates that reviewers with more eLife review experience tend to give more positive reviews. It is worth noting that there may be a selection bias in the invitation of reviewers if more positive reviewers are more likely to be invited again. To control for this factor, we studied reviewers who had reviewed 3 or more eLife submissions. As shown in Figure 3c, their average review rating in their first review for eLife was 0.441 ± 0.009, while in their second review it was 0.451 ± 0.009. In their third review, the average rating was 0.459 ± 0.009. These results indicate that new reviewers with less eLife review experience tend to give lower ratings.

### Analysis of reviewing editor decisions

The final step in the submission process is the decision made by the reviewing editors. Similar to the senior editors, the reviewing editors were also resistant to the busyness effect. Figure 4a illustrates the number of submissions decided upon each month and the average acceptance rate. Despite fluctuations in the number of submissions throughout the year, with peaks at the beginning of the calendar year and in the summer, the acceptance rate remains consistent throughout the year.

**Figure 4:**
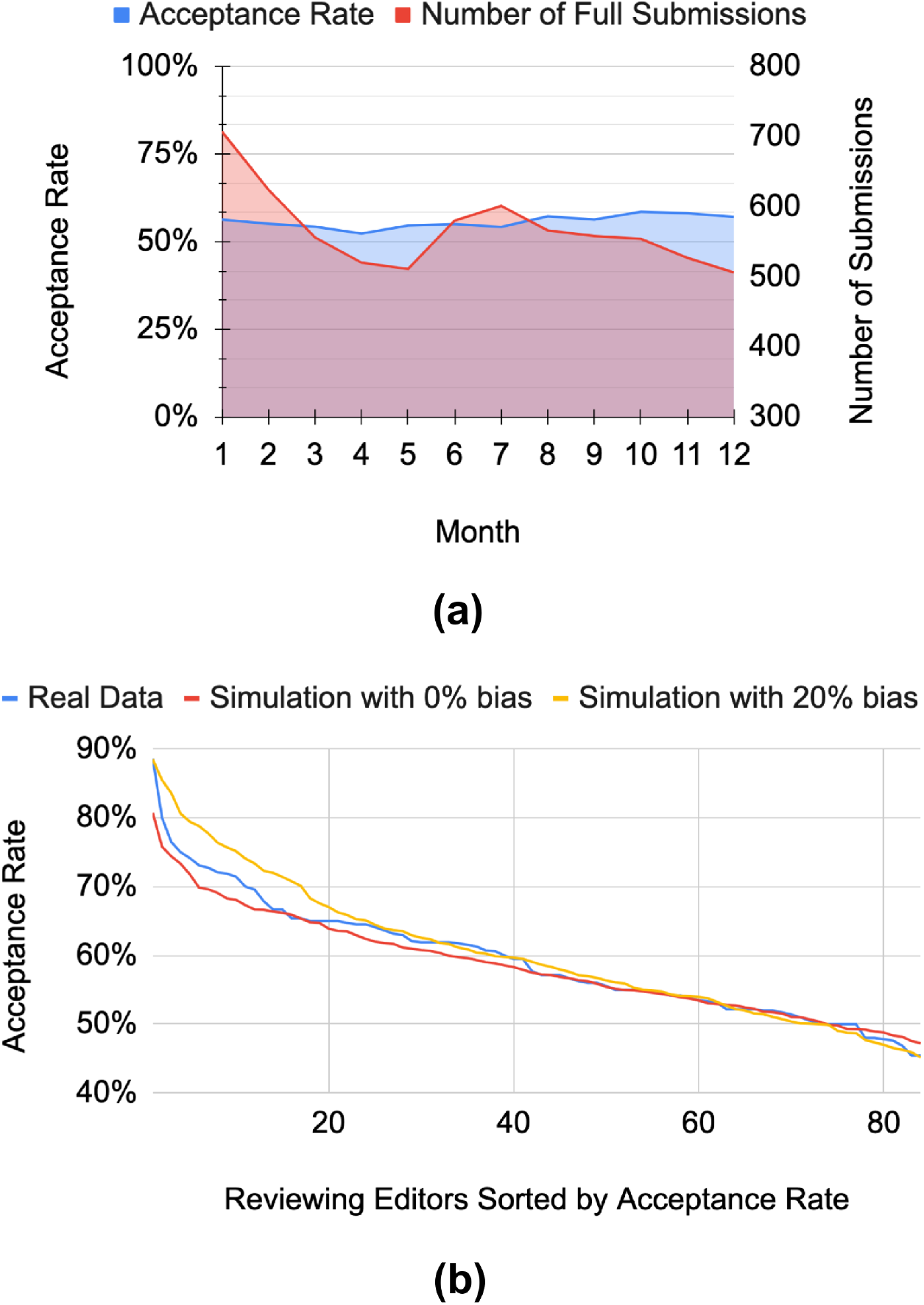
Analysis of reviewing editor decisions. (a) No busyness effect: The x-axis indicates the month of the decision, with the red curve representing the number of submissions and the blue curve representing the acceptance rate for that month. Although the number of submissions varies throughout the year, with peaks at the beginning of the calendar year and in the summer, the acceptance rate remains constant. (b) Heterogeneous acceptance rates: The y-axis represents the acceptance rate of each reviewing editor. The blue curve shows the actual distribution of acceptance rates among reviewing editors. The red and yellow curves represent simulations with 0% and 20% bias, respectively. If 20% of the reviewing editors were systematically biased as in our model, we would expect to see a difference between the curves. The lack of difference suggests that the amount of systematic bias is relatively small. It is worth noting that other bias models may be at play.

Similar to our simulation analysis of reviewer decisions, we sought to understand the role of bias in the acceptance rates of reviewing editors. To do so, we compared the real distribution of acceptance rates with simulated distributions representing different levels of bias. Figure 4b displays the results of our analysis. The blue curve represents the simulated distribution with 0% bias, in which each reviewing editor made random acceptance decisions by flipping a biased coin with probability p=0.54, which was the average acceptance rate of full submissions. This simulation represents a scenario in which all reviewing editors have an equal likelihood of accepting or rejecting a manuscript, regardless of any potential biases. The yellow curve represents the simulated distribution with 20% bias, in which 10% of the reviewing editors consistently gave higher ratings by flipping a biased coin with p=0.74, and another 10% consistently gave lower ratings by flipping a biased coin with p=0.34. This simulation serves as a power analysis to show that if 20% of the reviewing editors were systematically biased as in our model, we would have detected a difference in the curves. The fact that we did not observe such a difference suggests that the amount of systematic bias is relatively small. In addition to comparing the simulated and real distributions, we also analyzed the consistency of the acceptance decisions made by reviewing editors with the average ratings of the reviewers. We found a strong point biserial correlation coefficient of 0.592, which suggests that the reviewing editors largely follow the majority view of the reviewers. Overall, our analysis indicates that the heterogeneity of acceptance rates among reviewing editors can largely be attributed to randomness, and that the level of systematic bias is relatively low. While it is important to continue monitoring and addressing any potential biases, our findings suggest that the integrity of the review process is largely maintained.

Furthermore, our analysis did not reveal any significant differences in acceptance rates based on the gender or nationality of reviewing editors. There was no significant difference in acceptance rates between U.S. and non-U.S. reviewing editors (U.S. 0.560 ± 0.018 vs. non-U.S. 0.564 ± 0.016, χ2(df=1, n=6,498)=0.093, p=0.761) or between male and female reviewing editors (male 0.566 ± 0.015 vs. female 0.550 ± 0.024, χ2(df=1, n=6,498)=1.131, p=0.288). These findings suggest that the acceptance rates of reviewing editors are largely independent of these demographic factors.

## Discussion

In this study, we aimed to quantify the extent to which external factors, unrelated to the quality of a paper, affect its likelihood of acceptance in the peer review process. Understanding and reducing the influence of such random factors is important for improving the efficiency and fairness of the scientific ecosystem.

We studied two types of external factors that can introduce randomness: the timing of the submission, and the assignment of editors and reviewers. Overall, our study suggests that these factors have relatively small effects on the paper’s outcome. This supports the robustness of the peer-review process in *eLife*. One potential concern is the significant heterogeneity in the encouragement rates of different senior editors. While this may be partially due to the small number of senior editors and variations in the number and potential quality of submissions across disciplines, it is difficult to definitively determine whether this heterogeneity reflects any editor-specific biases.

Despite the insights gained from this study on the potential impact of external factors on the peer review process at eLife, there are several limitations to consider. One limitation is the lack of an objective measure of manuscript quality, as we relied on reviewer recommendations and final decisions as indicators rather than a more objective measure. Additionally, the observational nature of the study makes it challenging to establish a causal relationship between external factors like submission timing and review outcomes. While our sample size is large overall, it may not have sufficient statistical power to detect all possible relationships between review outcomes and external factors, such as the number of submissions processed by each reviewer or reviewing editor. However, we have taken steps to improve the robustness of our analysis, such as controlling for submission quality by studying the consistency of reviewer ratings within the same submission and conducting a power analysis to determine the sensitivity of our findings to potential bias among reviewers and reviewing editors.

Future research will be necessary to determine the extent to which the effects we observed in this study can be generalized to other peer review contexts. As an open-access journal, it is possible that the authors, editors, and reviewers at eLife may be more supportive of open access compared to the broader scientific community. Additionally, eLife has specific review policies, such as the consultation among reviewers, which may not be applicable to other peer review contexts. Therefore, additional studies in other journals would be an important direction of future work. Despite these limitations, this study provides valuable insights into the potential impact of external factors on peer review outcomes at eLife and serves as a starting point for future research on this topic.

## Code availability

Our analysis code is publicly available at: https://github.com/Weixin-Liang/eLife-peer-review-random-external-factors-study.

## Acknowledgement

We thank Andy Collings and James Gilbert from eLife for sharing the data and providing explanations, and eLife’s editorial leadership team for input and suggestions. This research is supported by NSF SCISIPBIO 2022435.

## Materials and Methods

### Details of the dataset

We were provided with metadata for research papers submitted to *eLife* between 2012 and 2018 for analysis. Our dataset contains the peer review outcomes of *eLife* from 2012 to 2018, with 34,161 initial submissions and 10,227 full submissions. As discussed in the results section, we focused on the period from 2016 to 2018, where the statistics of the number of submissions and selectivity were relatively stable, in order to ensure that our findings were robust. Therefore, we studied the peer review outcomes of 23,190 initial submissions and 6,498 full submissions to *eLife* that were submitted between 2016 and 2018. Each submitted manuscript was considered as our unit of analysis. For each submission, we had data on the full text, title, author name, author address, initial submission date, full submission date, all editorial decisions, and the time-stamp of the decisions. For each reviewer’s evaluation, we had the full evaluation text, predicted reviewer recommendation score (obtained through a machine learning approach, see below), and the internal reviewer identification information. For all individual analyses in our study, we analyzed senior editors who had reviewed at least 100 submissions, reviewing editors who had reviewed at least 20 submissions, and reviewers who had reviewed at least 10 submissions. For additional details on the *eLife* review process, we refer interested readers to Schekman et al. (2013).

Our *eLife* journal dataset was acquired through a data use agreement with the journal. All analyses of the data followed research guidelines for the use of high-risk data on human subjects (University IRB protocol 12996). In full compliance with the signed protocol, data were stored in the Stanford Neuro Computing Platform, which is specifically designed for High-Risk Data.

### Quantifying reviewer recommendations with machine learning

Since *eLife* reviews do not include an explicit review rating score, we performed sentiment analysis on the review text to quantify its positivity. To quantify reviewers’ recommendations to the editor for a submitted manuscript, we first fine-tuned a BERT deep neural language model (Devlin et al., 2019) using a set of openly available reviews from the PeerRead corpus (Kang et al., 2018). We then fine-tuned these reviews on a training set of 900 *eLife* reviews that were manually scored on a 4-point scale (reject/mixed/accept/strong accept). Using a held-out test set, we found a high correlation (r = 0.78) between the hand-scored reviews and the automatically scored reviews. Importantly, accuracy was 100% for reviews hand-scored at extreme values - i.e., there were no reviews hand-scored as a 1 (“reject”) that were automatically scored above 3.0 or *vice versa*. The deep neural language model was implemented using PyTorch 1.4.0.

In addition, since there are usually multiple reviews (usually three) for the same submission, we controlled for submission quality by studying whether some reviewers consistently have higher or lower review ratings compared to other reviewers of the same submission. To do this, we studied the rank of each review’s rating among the reviews for the same submission. We normalized the rank by the number of reviews for the same submission. For example, for a submission with 3 reviews, the most negative reviewer would have a normalized rank of 0.0, the second most negative reviewer would have a normalized rank of 0.5, and the most positive reviewer would have a normalized rank of 1.0. We used the average review rating rank to study the heterogeneity of reviewers.

### Statistical analysis

Data processing, statistical testing, and visualization were performed using Python version 3.7. We used the Python package “sklearn” (https://scikit-learn.org/) to conduct a series of chi-squared tests of equal proportion. P-values and confidence intervals are provided as a means of interpretation; we follow the convention of using a threshold of 0.05 for statistical significance. When visualizing proportions in figures, 95% confidence intervals are calculated using the formula 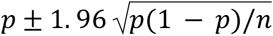, where p is the proportion and n is the number of observations in the group. We performed t-tests to compare the means of two groups. We used the Python package “matplotlib” (https://matplotlib.org/stable/index.html) to generate our plots. In accordance with the data-sharing protocol, we have carefully avoided reporting any identification information for editors or reviewers to protect their identities.

### Gender and nationality analysis

We follow the method used by previous research (Larivière et al., 2013; Helmer et al., 2017) to identify the gender of reviewers and reviewing editors based on their names. The method has recently been validated on a dataset of scientist names extracted from the WoS database (Santamaría & Mihaljević, 2019). We used the Python package “gender-guesser” (https://pypi.python.org/pypi/gender-guesser/) to infer the gender of each individual based on their name. According to Santamaría and Mihaljević (2019), the “gender-guesser” package has the lowest misclassification rate and minimizes bias, with a misclassification rate of 1.5% for European names, 3.6% for African names, and 6.4% for Asian names. We used the provided affiliation information to match the nationality of each individual.

## Supplementary Figures

**Figure S1:**
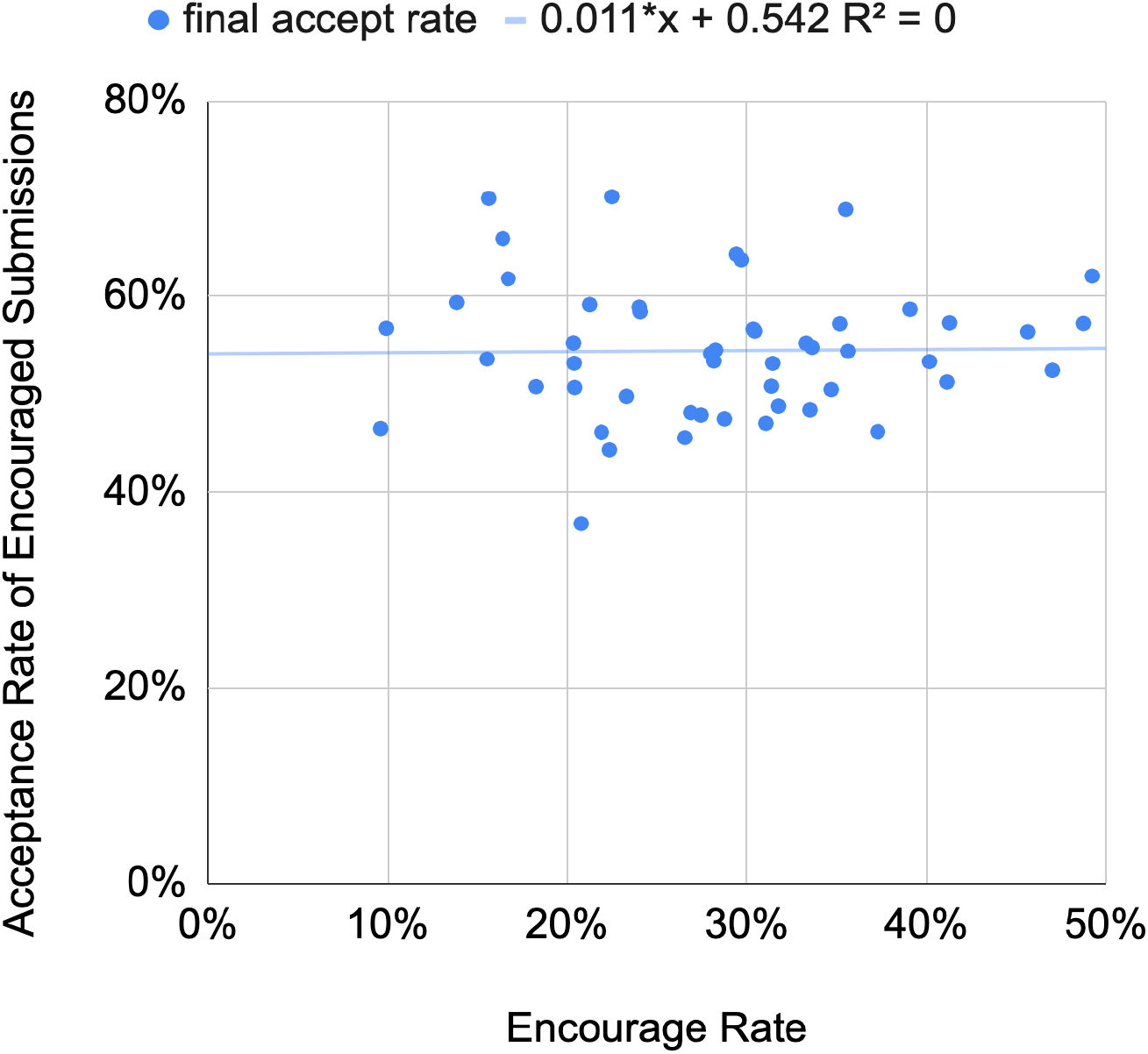
Encouragement rate is not correlated with the acceptance rate of encouraged submissions. Each blue dot represents a senior editor. The x-axis represents the senior editor’s encouragement rate, and the y-axis represents the acceptance rate of submissions that were encouraged by the senior editor. Our analysis does not reveal a significant correlation between senior editors’ encouragement rates and the acceptance rate of the submissions they encourage.

**Figure S2:**
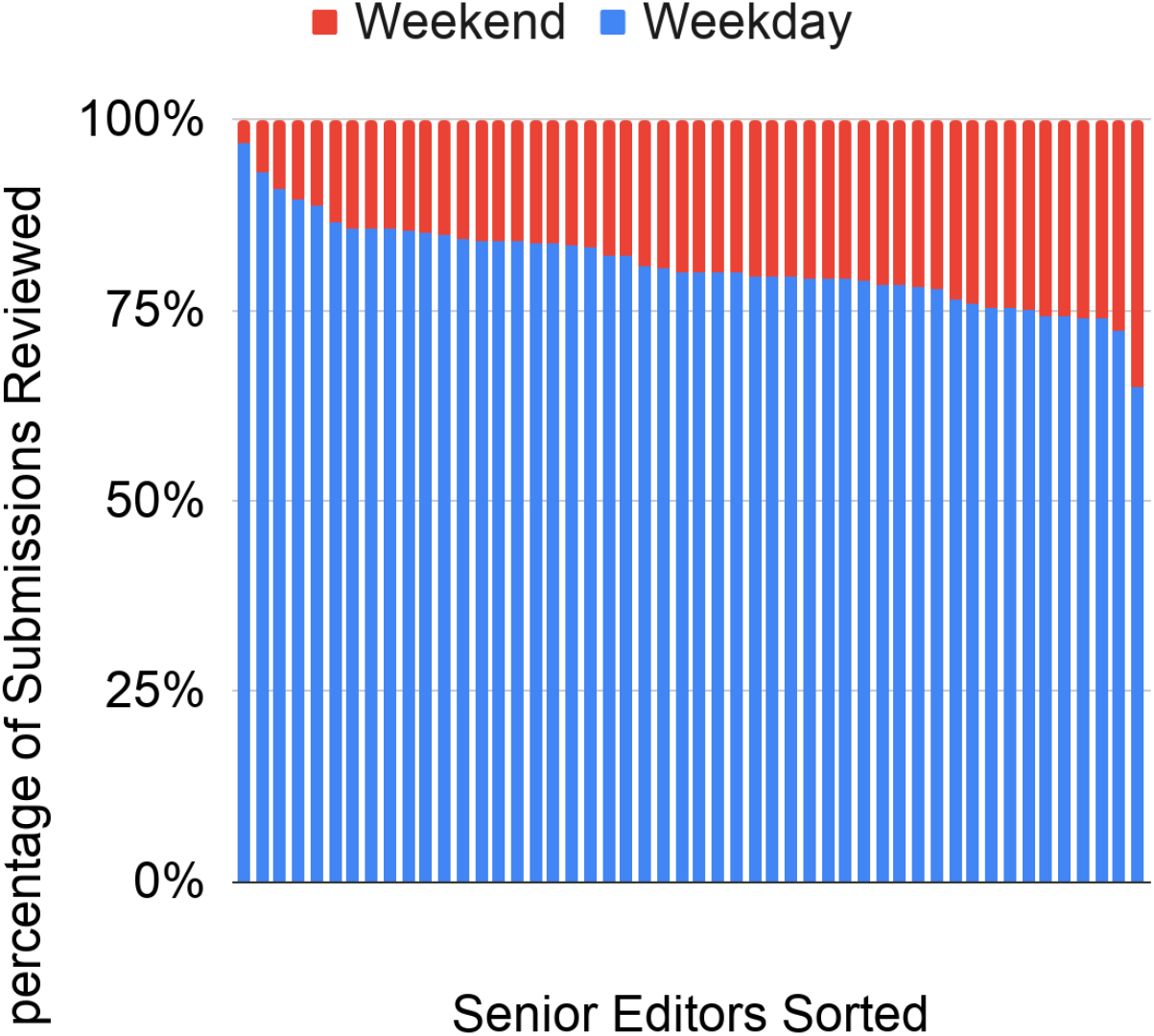
Comparison of the number of initial submissions reviewed on weekdays and weekends by senior editors. The x-axis indicates the senior editors, and the y-axis represents the normalized count of submissions. On average, senior editors review 76% of the submissions during weekdays and 24% of the submissions during weekends. Furthermore, none of the senior editors review more submissions on weekends than on weekdays.

**Figure S3:**
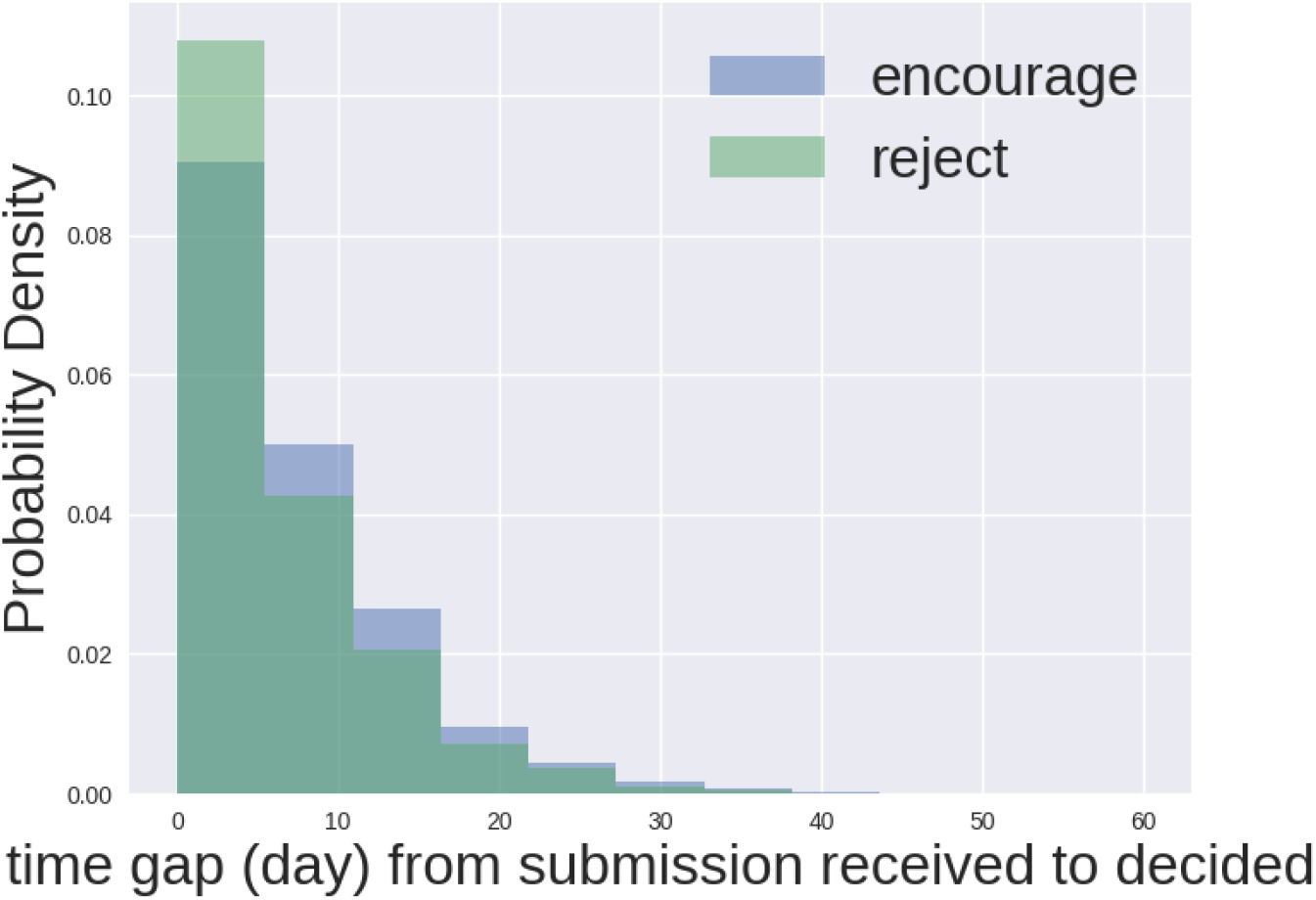
Rejection decisions by senior editors are faster on average. On average, it takes 7.34 days for a senior editor to make an encourage decision for an initial submission and 6.09 days for a senior editor to make a reject decision. One possible explanation is that senior editors may reject initial submissions without consulting reviewing editors, while reviewing editors are almost always consulted for encourage decisions. This is because the senior editor assigns the reviewing editor for an encourage decision in *eLife*. Therefore, some reject decisions may have one less step and be faster on average.

